# Simultaneous analysis of single-cell gene expression and morphology provides new insight into how microglia change with age

**DOI:** 10.1101/2025.02.28.640888

**Authors:** Douglas E. Henze, Andy P. Tsai, Tony Wyss-Coray, Stephen R. Quake

**Affiliations:** Department of Bioengineering, Stanford University, Stanford, CA, USA; Department of Neuroalogy and Neurological Sciences, Stanford University School of Medicine, Stanford, CA, USA; Wu Tsai Neurosciences Institute, Stanford University, Stanford, CA, USA; The Phil and Penny Knight Initiative for Brain Resilience, Stanford University, Stanford, CA, USA; Department of Applied Physics, Stanford University, Stanford, CA, USA; Chan Zuckerberg Initiative, Redwood City, CA, USA

## Abstract

Cellular morphology is intimately connected with function. While the link between morphology and functional states has been studied extensively, the role of subcellular transcript localization in cellular function remains unclear. Here we use microglia, the brain’s resident macrophages, as a model to dissect the interaction of morphology, transcript localization, and function. Using multiplexed error-robust fluorescence in situ hybridization combined with fluorescent immunohistochemistry, we analyzed transcript distribution and morphology simultaneously in young and aged mouse brains. Our approach revealed how mRNA spatial organization varies across microglial states. We identified distinct transcript localization patterns within microglial processes and uncovered morphological heterogeneity within transcriptomically defined populations. Notably, we found a subpopulation of disease-associated microglia with a ramified morphology (displaying numerous processes), challenging the conventional assumption between morphology and microglial states. Finally, we found that aging not only alters the distribution of compartmentalized mRNAs but also reshapes their colocalization networks, shifting microglial functions from synaptic maintenance and phagocytic processes in younger brains to migration and catabolic pathways in older brains. Our findings highlight the role of subcellular transcript organization in shaping microglial morphology and function, offering new avenues for studying and modulating microglial states in health, disease, and aging.

## Main

Subcellular localization of mRNA provides precise spatial control over protein synthesis^1^. This process is essential for cells with complex morphologies, and its disruption has been linked to various age-related diseases^2–8^. In the central nervous system (CNS), neurons and glia rely on localized transcripts to maintain polarized functions^9–13^. Microglia—the resident immune cells of the CNS^14^— are an ideal model for studying how mRNA localization gives rise to functional diversity through interaction with cellular morphology^15,16^. This is because traditionally microglia have been classified into two morphological states with distinct functionalities: a “homeostatic” functionality with ramified or branched processes, and an “activated” (disease-associated) functionality with an amoeboid shape^17^. However, the mapping between morphology and function can be more complex than this simple picture might suggest, as recent findings showed that ramified microglia can perform phagocytic functions typically associated with amoeboid morphology^18^.

Single-cell transcriptomics has been used to characterize myriad microglial states^19–21^— including homeostatic^22^, transitioning^23^, and disease-associated microglia (DAM) states^24^—across healthy and diseased brains^25,26^. However, that work relied on tissue dissociation, which obscures spatial context of cells and fails to capture cellular morphology or subcellular transcript localization. In contrast, spatial transcriptomics preserves tissue architecture, but most approaches lack the resolution needed to analyze mRNA distribution at the subcellular level^27^. Techniques like multiplexed error-robust fluorescence in situ hybridization (MERFISH) address this limitation, offering high-resolution mapping of mRNA and enabling studies of aging and disease in the CNS^28–30^. Although a recent study used spatial transcriptomics and electron microscopy on adjacent sections to correlate microglial ultrastructure with the transcriptome^31^, no current method captures subcellular transcript location along the full complexity of their processes. Some evidence suggests that mRNA positioning in microglial processes is implicated in cell function^32^.

To address this, we combined MERFISH with high-content fluorescence immunostaining to explore how transcript localization correlates with microglial morphology in young (3-month-old) and aged (24-month-old) mouse brains. Our analysis uncovered a diverse array of microglial morphologies associated with various transcriptomic phenotypes, including a distinct subpopulation of DAMs characterized by highly ramified processes. By correlating transcriptomic data with morphological features, we identified genes strongly associated with this cell state, potentially impacting microglial function. Additionally, spatial mapping of transcripts within individual microglia revealed a set of process-localized genes predictive of specific morphological states across the lifespan. Finally, we found that aging not only alters the distribution of compartmentalized mRNAs but also reshapes their colocalization networks, shifting microglial functions from synaptic maintenance and phagocytic processes in younger brains to migration and catabolic pathways in older brains. Collectively, these findings demonstrate that subcellular mRNA organization—both in terms of localization and age-dependent clustering—helps define distinct morphological phenotypes and functional states of microglia, thereby providing new insights into how localized transcriptomic architecture influences microglial roles in the aging brain.

## Results

### Microglial transcriptional state depends on age and region

To visualize gene expression and image microglial morphology *in situ,* we applied subcellular resolution spatial transcriptomics using MERFISH combined with fluorescent immunostaining of ionized calcium binding adaptor molecule 1 (IBA1)^33^ on sagittal sections of six mouse brains for each sex at two ages: 3 months (young) and 24 months (aged) (**Figure 1A**, **Methods**). Our spatial transcriptomics data contains count and subcellular localization information for a panel of 500 curated genes, designed to capture cell type and age-related variation across the brain (**Figure 1B**, **Methods**). This gene panel was used to segment^34^ and characterize spatially resolved single-cell gene expression. After quality control filtering (**Extended Data Fig. 1A & 1B**), preprocessing, and clustering of spatially resolved single-cell transcriptomes, we identified and labeled^28^ 991,315 cells across 35 major cell classes, distributed across seven major brain regions (**Methods, Extended Data Fig. 1 C-E**).

**Figure 1:**
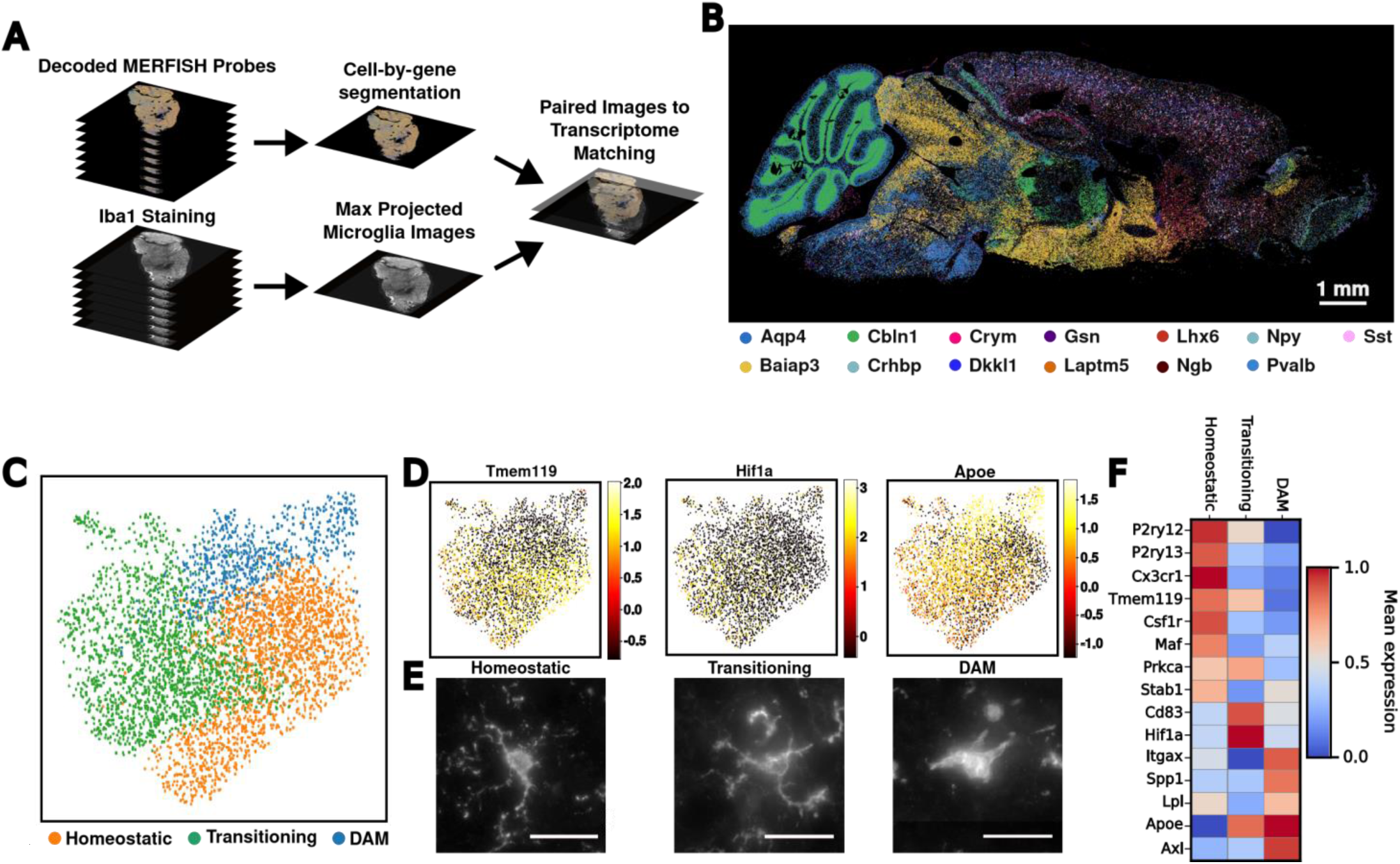
MERFISH paired with fluorescent IHC allows for single microglia transcriptomics and imaging. **A,** Schematic overview of the image processing pipeline to generate max projected transcriptome decodings and images of microglia across the whole mouse brain. **B,** Neuronal and glial markers resolved in space. **C,** UMAP plots of microglia colored by identified clusters. **D,** UMAP plots of microglia colored by marker gene expression. **E,** Representative images of IBA1 stained microglia from each of the identified clusters. Scale bars are 25 µm. **F,** Heatmap of average expression for each microglia cluster.

To characterize microglia’s regional distribution and transcriptional states in the aging mouse brain, we identified and reanalyzed microglia across the 12 brain sections and across two age groups (**Methods**). After identifying transcriptomically defined microglia, we aligned the transcriptomics data to the fluorescent imaging data and used the segmentation of the IBA1 channel to refine microglial cell boundaries (**Extended Data Fig. 2A-C**). We then recomputed the number of RNA molecules that localized within the microglial cells to generate single-cell gene expression counts for microglial cells alone (**Extended Data Fig. 2D**). In total, we identified 4,465 microglial cells (**Methods**) which could be directly aligned with the IBA1 segmentation representing their morphologies. The microglia were predominantly located toward the posterior brain, specifically in the midbrain, pons, and medulla (MPM) regions (**Extended Data Fig. 3A**). Clustering analysis of these aligned microglia visualized on a 2D uniform manifold approximation and projection (UMAP) embedding^35^ revealed three primary subtypes representing microglial transcriptional states (**Figure 1C-E, Extended Data Fig. 3B**). The homeostatic cluster, marked by the expression of canonical microglial markers *P2ry12*, *Tmem119*, and *Cx3cr1*^22^, was enriched in the cortex (F**igure 1D**; **Extended Data Fig. 3C**). The most prevalent cluster, identified as transitioning microglia, expressed high levels of *Cd83* and *Hif1a*^23^ and was primarily localized to the MPM and the hypothalamus (**Figure 1D**; **Extended Data Fig. 3D**). The third cluster, defined by elevated expression of DAM markers *Itgax*, *Axl*, and *Spp1*^24^, was primarily found in the cerebellum, hippocampus, and MPM (**Figure 1D**; **Extended Data Fig. 3E**). DAM was the main microglial subtype which was increased in aged mice as compared to young mice (**Extended Data Fig. 3B**). There was a small, region-dependent effect in which the transcriptomes of microglia in subsets of the cortical layer and cerebellum deviated slightly from that of the other populations (**Extended Data Fig. 3F**).

Given that microglial function is typically associated with morphological changes, we next aimed to quantify morphological differences among the various transcriptomic classes. Using the same IBA1 segmentation, we calculated 32 morphological features for all microglia and compared them between transcriptomic classes (**Extended Data Fig. 2C; Extended Data Fig. 4**). As expected, DAM exhibited a more amoeboid morphology, with elevated values of cell solidity (**Extended Data Fig. 4**) compared to the other two classes on average^36^. Interestingly, both transitioning and homeostatic microglial clusters showed considerable variation in the grading of their ramification (displaying numerous processes) status, with minimal differences in the number of branching points in the processes. Furthermore, we observed high variance in the morphological characterization of DAMs, with some DAMs displaying ramification properties similar to homeostatic microglia (**Extended Data Fig. 4**). These findings suggest a disconnect between transcriptomic classification of microglia and their inferred morphological ramification patterns and point to an additional heterogeneity in the function of DAMs based on their morphological properties.

### Microglial functional states defined by transcriptomic states have heterogeneous morphology

After examining the morphologies associated with the different transcriptomic states of microglia, we aimed to determine whether their underlying geometry could provide deeper insight into the cells’ functional states. To quantify microglial morphology, we used the previously aligned maximum projection images across all planes of the IBA1-stained sections to the Bayesian-defined cell boundary priors to generate single-cell microglial images (**Extended Data Fig. 5A**). We then used a pre-trained neural network to parameterize these images and create a high-dimensional embedding of the morphological space^37^ (**Extended Data Fig. 2B**). We then clustered the microglial cells using the microglial morphology space and visualized them on a UMAP embedding (**Figure 2A**). Applying k-means clustering to microglial cells we observed five clusters of microglial cells in this morphological space along a continuum from the least to the most ramified morphologies (**Figure 2B**). The clusters were named along the morphological trajectory from C1-5 with the C1 cluster having the lowest values of ramification features such as cell area, the number of terminal points, and the fractal dimension, and the C5 cluster possessing the highest values. The regional distribution of microglial shapes is consistent with previous studies, with more ramified microglia primarily located in the cortex, while amoeboid microglia were predominant in the mid- and hindbrain^38^ (**Extended Data Fig. 5B-F**). As previously observed, the number of amoeboid microglia increased with age (**Extended Data Fig. 5G**)^39^. The morphological classes were characterized by variations in both basic measurements such as cell area and solidity, as well as established ramification metrics like fractal dimension and the number of terminal branches^40–43^ (**Figure 2B**). These measurements confirm that the morphology embedding space effectively captured the variation in microglial ramification (**Extended Data Fig. 6**).

**Figure 2.**
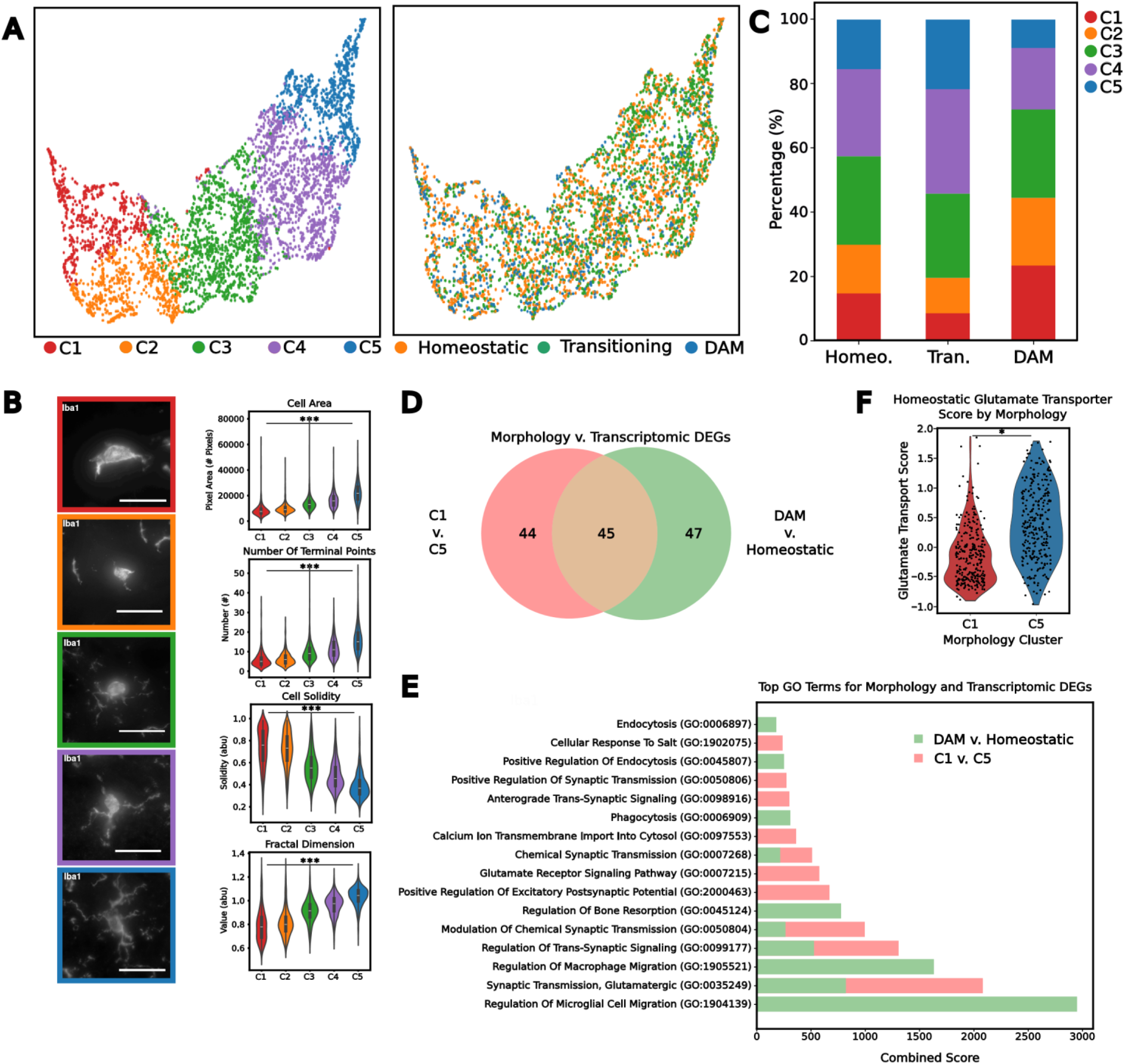
Morphologically Defined Microglia are Functionally Distinct from Transcriptomically Defined Microglia. **A,** Morphological UMAP embeddings of microglia, colored by leiden clusters (left) and by identified transcriptomic clusters (right). **B,** Representative images of microglia from each of the morphologically defined leiden clusters (left) and a selection of manually measured morphological features for each leiden cluster (right). Scale bars are 25 µm. **C,** Frequency of morphological leiden clusters per microglia transcriptomic cluster. **D,** Comparison of differentially expressed genes between C1 and C5 and between DAM and Homeostatic microglia. **E,** Gene ontology biological processes terms, sorted by combined score, for each of the comparisons made in D. **F,** Comparison of a glutamate transporters scored expression between different morphologically classified cells amongst homeostatic microglia. The * denotes a p-value < 0.05, and *** denotes p-value < 0.001.

We calculated the percentage composition of different morphological classes within each transcriptomic classification to evaluate the overlap between the morphology-derived microglia clusters and the transcriptomically-derived clusters (**Figure 2A & C**). As expected, the transcriptional state of microglia was associated with reduced ramification, evident in the increased proportions of morphology classes C1 and C2, which represent an amoeboid morphology, and the lower percentages of classes C4 and C5, which represent a ramified morphology, in the DAM cluster. C3 morphological cluster of microglial cells were uniformly distributed across all transcriptomics classes. Interestingly, we find a small proportion of homeostatic microglia displaying amoeboid morphology as well as a novel population of DAM cells displaying ramified morphology (**Figure 2C**). These ramified DAMs representing ∼ 25% of the transcriptomic class might be involved in actively patrolling the CNS while maintaining their phagocytic and pro-inflammatory function. Additionally, we see that transitioning microglia displayed the highest proportion of ramified morphologies, suggesting that homeostatic microglial cells display a slight increase followed by a decrease in ramification when undergoing functional responses (**Figure 2C**). These findings demonstrate the presence of a morphological gradient amongst the different microglial transcriptomic classes. Among microglia with distinct transcriptomic states, the striking morphological heterogeneity supports the now accepted concept that function, morphology, and transcriptional state are not necessarily connected^44^. The variable proportions of the different morphologies for each transcriptomic class also imply a range of the functional implications that the morphological states can have on the different transcriptomic classes, further skewing previous definitions of microglial cell state.

To further explore the association between transcriptional state and morphology, we performed differential gene expression analysis between the morphologically defined clusters C1 and C5, which represent the two extremes of the microglial morphology space, as well as between the transcriptomically defined homeostatic and DAM clusters, the two extremes of the microglial reactive states (**Figure 2D**). Comparative analysis of the most enriched Gene Ontology (GO) biological process terms derived from these differentially expressed gene (DEG) sets showed that the transcriptomic changes between homeostatic and DAM microglia were associated with ontology terms such as cell migration and phagocytosis. In contrast, DEGs between the two extremes of the morphology space were linked to synaptic signaling and transmembrane transport (**Figure 2D & 2E**), suggesting functional differences between cells of varying geometries that are independent of changes between homeostatic and DAM phenotypes. Together, these findings indicate that reduced ramification does not necessarily correlate with the DAM functional state. Additionally, comparing the morphologically distinct cells within the homeostatic microglia, we observed that genes related to glutamate trafficking and to the anchoring of glutamate receptors, which are associated with DAM microglia (**Methods)**, are elevated in the ramified microglia as opposed to the amoeboid-shaped cells (**Figure 2F**). These results contradict that increased expression of different glutamate trafficking genes results in activated and morphologically less complex cells as observed previously^45,46^. Collectively, comparing our morphology-based clustering to the transcriptomics-based classification of microglia revealed previously unrecognized heterogeneity, which could not be fully explained through either transcriptomic or morphological analyses alone.

### Gene-morphology correlations explain shape heterogeneity

Given the observed disconnect between transcriptomic states and cell morphology, we sought to investigate potential dependencies between morphological features and gene expression. Previous research has suggested a relationship between microglial gene expression and ultrastructural features^31^. To determine whether similar dependencies exist between cellular morphological features and the transcriptome, we performed a Spearman correlation analysis between gene expression and morphological features using our paired morphological and transcriptomic measurements for the microglia across all brain regions (**Figure 3A-C**). Correlation analysis revealed two distinct clusters of genes and morphological features. The morphological features associated with a more amoeboid cell shape correlated with genes such as *Csf1r*, *Cx3cr1*, *Csf3r*, and *Mrc1*. In contrast, genes such as *Slc1a2*, *Gria2*, *Pink1*, and *Ank2* correlated with ramification features. These two distinct clusters identify potential genes which can be used as markers for the morphological status of microglia.

**Figure 3.**
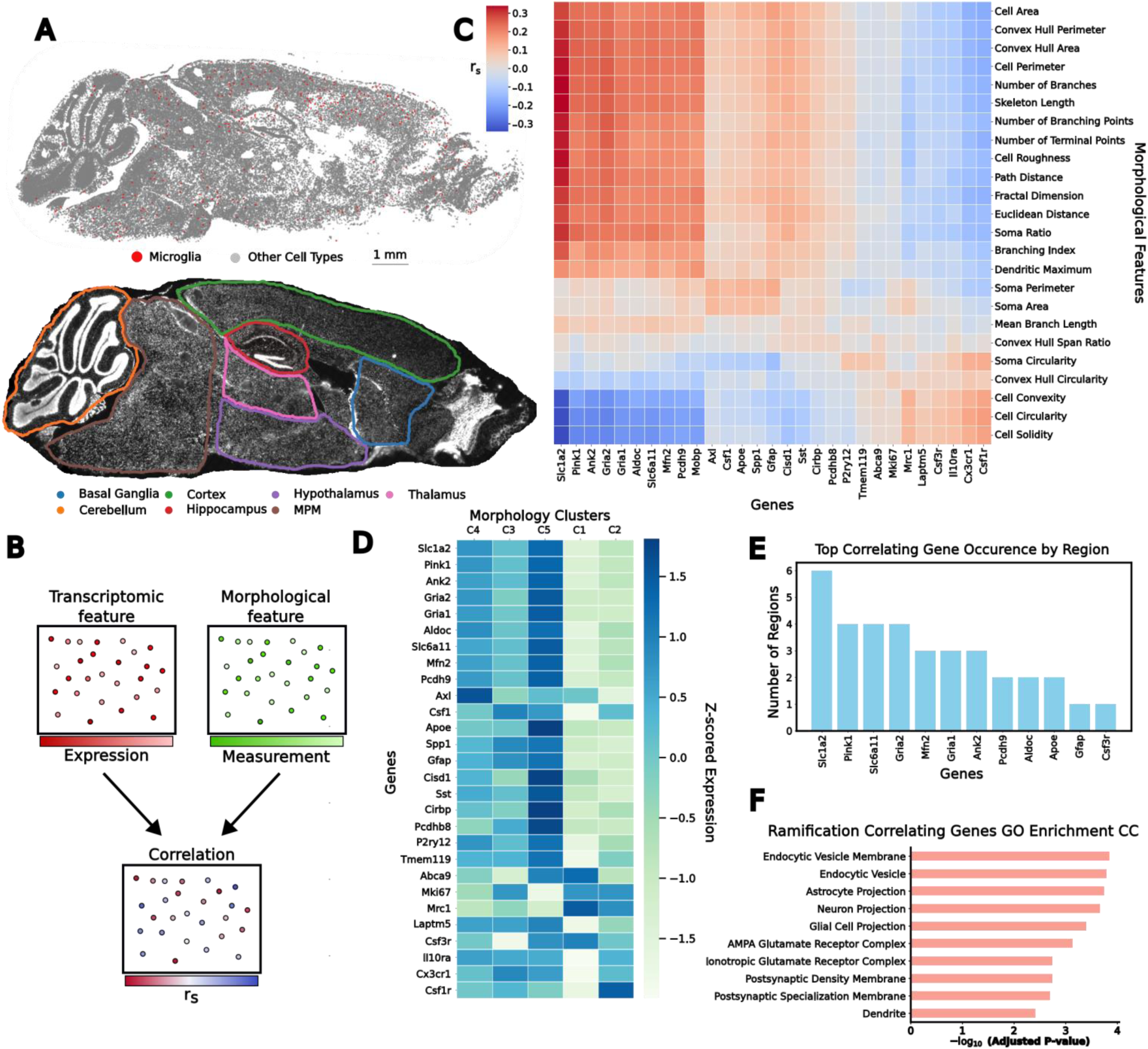
Morphology-Transcriptome Correlation Identifies Compartment Specific Genes. **A,** Schematic image of microglia amongst all other cells in the brain (top) and image of the same brain separated into the brain regions (bottom). **B,** Schematic representation of workflow for establishing correlation results, for every transcriptomic feature we use a morphological feature on the same microglia to calculate the Spearman correlation. **C,** Clustered heatmap showing the Spearman correlation between different microglial morphological features and select genes. **D,** Heatmap showing the ability of select genes to be able to cluster the cells morphological classifications **E,** Genes amongst the top correlating genes for morphology characteristics in each region of the brain, sorted by number of regions in which they occur. **F,** Gene ontology cell compartments terms for the genes which are amongst the top correlating genes in any segmented region of the brain.

Using the transcriptomic features of morphologically distinct microglial cells, we next sought to curate a gene set that could explain variance in microglial cells in both morphological and transcriptomics space. We combined the top five genes correlated to each morphological property to create a gene set of 28 morphology-associated genes. Notably, clustering of an additional held-out population of cells using this gene set separated the cells into two distinct groups, containing distinct morphology clusters defined previously **(Figure 3D**). Clusters C3-C5 corresponded to genes associated with ramified morphology, while clusters C1 and C2 were associated with genes linked to a more amoeboid morphology **(Figure 3D**). These results clearly identify a set of genes which can be used to cluster microglial cells with distinct morphological properties.

We also observed that individual morphological measurements correlated with genes that are functionally relevant to the compartment from which the measurement was derived (**Extended Data Fig. 7A-C)**. For instance, fractal analysis, which is commonly used to assess microglial complexity^47^, largely relates to the complexity and branching of microglial projections. Genes with the highest correlation to a cell’s fractal dimension were enriched for cell compartment ontology terms related to morphological projections (**Extended Data Fig. 7A**). Similarly, genes with highest correlation to soma area, which is a characteristic of activated microglia that play an active role in inflammation^48^, were enriched for organelle-related ontology terms that are consistent with delivery of cell signaling molecules, particularly those involving the endoplasmic reticulum (**Extended Data Fig. 7B**). Interestingly, cell solidity—a metric canonically associated with amoeboid shape and therefore phagocytotic potential^49^–was correlated with genes enriched for GO terms involving primary lysosomes and endocytic vesicles (**Extended Data Fig. 7C**).

Since the brain exhibits regional heterogeneity in response to inflammatory stimuli^50,51^, we next investigated whether regional differences influenced the relationship between microglial gene expression and ramification features. For each of the seven brain regions quantified in our sagittal section MERFISH dataset (**Figure 3A**), we calculated the Spearman correlation for the genes associated with the ramification features (**Methods**). From these, we compiled a list of the top five genes with the highest average Spearman correlation across all ramification features in each brain region (**Figure 3E**). Only *Slc1a2*, which codes for the glutamate transporter 1 (GLT-1) protein, was among the top ramification correlated genes in more than 5 brain regions. The genes included in this region-specific correlation analysis were also related to functional compartments of the cell. Notably, the genes which correspond to ramification features across the brain correspond to GO terms related to glial cell projections (**Figure 3F**). Collectively, these findings suggest that the expression of genes linked to different subcellular compartments can effectively cluster cells morphologically.

### Process enriched transcripts are a predictor of morphology

The subcellular localization of mRNAs is known to influence functional differences in both neuronal^6^ and glial cell types^52^. Because genes that correlate strongly with microglial ramification features correspond to GO terms related to glial cell projections, we investigated how the distribution of transcripts across distinct subcellular compartments might relate to morphological states. To do this, we quantified the normalized distance of each transcript from the soma center in the two most ramified microglial clusters (C4 & C5) (**Methods**) (**Figure 4A**). Our analysis showed that homeostatic microglial marker genes such as *Cx3cr1*, *Csf1r*, and *Laptm5* were enriched closer to the soma, whereas genes like *Apoe* and *Slc1a2* displayed a more uniform distribution from the nucleus out toward the periphery of the processes. This suggests a compartment-specific organization of mRNAs within ramified microglia. To more thoroughly examine this compartmentalization, we computationally segmented microglia into soma and process compartments, then quantified the RNA abundance of all 500 measured genes in each subcellular region (**Extended Data Fig. 2D**; **Figure 4B**).

**Figure 4.**
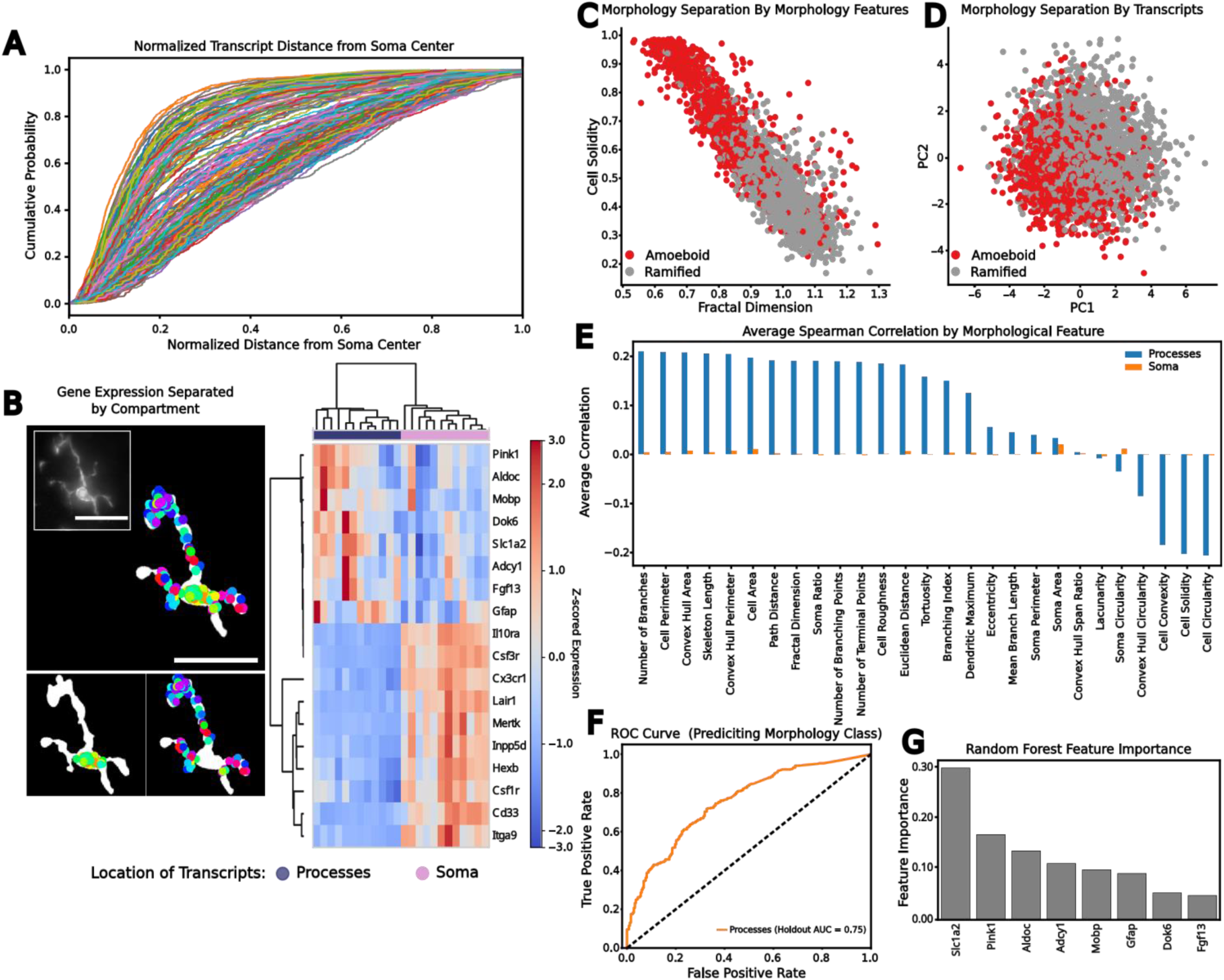
Compartment Specific Genes Can Predict Cell Morphology. **A,** The cumulative probability of transcript distance to the soma, normalized for maximum distance within each C4 and C5 cell. Different colored lines represent different genes. **B**, Schematic demonstrating the separation of microglia into soma counts and process counts (left) and clustermap showing genes cluster by location of transcripts(right). Scale bars are 25 µm. **C,** Amoeboid and ramified cells separate based on morphological features. **D**, Amoeboid and ramified cells separate based on the first and second principal component of compartment enriched genes. **E**, Average correlation between branch and soma enriched genes with morphological characteristics, colored by compartment genes were enriched in. **F**, Morphological classifier performance as quantified by receiver operator characteristic curve (ROC). **G,** Importance of individual genes in classifier as calculated through mean decrease in impurity.

We next compared the expression of transcripts localized to the soma versus those enriched in processes within C4 and C5 clusters to identify genes potentially involved in glial cell projections (**Methods**). From this analysis, we found 76 soma-enriched and 8 process-enriched genes (**Extended Data Fig. 8A; Supplementary Sheet 1**), which clustered based on their compartment-specific expression (**Figure 4B**). Gene set enrichment analysis revealed that soma-enriched genes were associated with terms related to lysosomes and phagocytic vesicles—consistent with their expected subcellular localization—while process-enriched genes were linked to extensions and branch-like structures present in both glial cells (**Extended Data Fig. 8A–C**). Together, this data supports the idea that distinct sets of genes localize to the soma or processes, potentially enabling compartment-specific functions.

Recognizing that genes correlated with ramified morphology encode terms reflecting projection cellular compartments, we hypothesized that compartment-enriched gene expression might serve to distinguish microglial morphology states. To test this, we separated the microglial population into ramified (C4 and C5) and amoeboid (C1 and C2) subpopulations (**Figure 4C**). We then performed principal component analysis (PCA) on the subset of compartment-enriched genes (those localized to the soma or processes) across the C4 and C5 cells. Using whole-cell log-normalized counts from both ramified and amoeboid microglia, the first two principal components provided modest separation of these morphological types (**Figure 4D**). To further assess whether compartment-enriched genes could classify microglial morphology, we computed the average Spearman correlation between each compartment’s gene set and morphological features. Process-enriched genes exhibited a stronger correlation with morphological traits than soma-enriched genes (**Figure 4E**), suggesting that the expression of process-localized transcripts may better define the morphological phenotype.

To evaluate the predictive power of process-enriched genes, we constructed a random forest classifier using their whole-cell log-normalized expression to distinguish ramified from amoeboid microglia. Training on 80% of the microglial cells and evaluating on a held-out set, the classifier achieved a reasonable accuracy, with an AUROC value of 0.75 (**Figure 4F**). Examining the genes most important for classification revealed *Slc1a2* and *Pink1* as the top contributors (**Figure 4G**), consistent with previous findings that both genes correlate positively with a more “activated” microglial state and function^45,53^. These results indicate that the transcriptional profile of a microglial cell is closely linked to the ramification of its processes and that the subcellular distribution of transcripts plays a pivotal role in shaping microglial morphological phenotypes. By revealing how compartment-specific RNA localization influences cell state, these findings support the notion that subcellular transcript distribution can differentially impact microglial functions depending on where those transcripts reside.

### Aging disrupts compartmentalization of transcripts

The compartmentalization of cellular functions within the brain is thought to be influenced by aging^54^. Since process-localized transcripts can effectively predict microglial morphology, we next examined how age affects transcript localization and the capacity to predict morphological states based on these compartmentalized gene sets.

To assess the impact of age, we compared soma- and process-enriched transcripts in the two most ramified microglial clusters at both young and aged time points (**Methods**). In young microglia, we identified 67 soma-enriched genes and 18 process-enriched genes. In contrast, in aged microglia we found 60 soma-enriched and only 5 process-enriched genes—both sets effectively clustered according to compartment-derived gene expression (**Extended Data Fig. 9A, B; Supplementary Sheet 1**). The reduced number of both soma- and process-enriched genes in aged cells suggests a diminished compartmentalization of mRNAs in older microglia.

We next evaluated whether these process-enriched genes maintained their predictive value for morphology across different ages. Using the previously identified process-localized gene sets (**Extended Data Fig. 8A**), we calculated the average Spearman correlation coefficient between these genes and morphological features in both young and aged microglia. We observed that process-enriched genes in young microglia correlated more strongly with morphological traits than the same gene set in aged cells (**Extended Data Fig. 9C**). To further test the predictive capacity of process-enriched genes across age, we trained random forest classifiers using the process-enriched gene expression profiles in young and aged microglia, each model trained on an 80% random sample of that age group. The resulting models achieved AUROC values of 0.76 for young microglia and 0.78 for aged microglia, reflecting stability in predictive power with age (**Extended Data Fig. 9D**).

Finally, we examined the key drivers of classification across ages by calculating the relative importance of each gene in the classifiers. This analysis again highlighted *Slc1a2* and *Pink1* as principal factors distinguishing morphological states in both age groups (**Extended Data Fig. 9E**). Taken together, these findings suggest that while aging may disrupt the extent of RNA compartmentalization, the core set of process-enriched, DAM-related genes like *Slc1a2* and *Pink1* remain integral to classifying microglial morphology.

### Subcellular transcript co-localization changes with age

The compartmentalization of mRNA, and its resulting functional impact, appears to shift with age. Previous work has shown that functional differences arising from mRNA compartmentalization are influenced not only by the localization of transcripts but also by their spatial proximity to one another^55–57^. Given that aging has been shown to affect mRNA compartmentalization within the brain^58–60^, we next examined how it influences the spatial clustering of mRNAs within specific subcellular compartments. To quantify clustering, we applied a generalized Ripley’s K analysis, a method designed to assess transcript clustering against a spatially randomized distribution^61–63^ (**Methods**). Since we had observed that expression patterns vary between soma and processes with age, we conducted separate clustering analyses for each compartment in the C4 and C5 cells (**Figure 5A**). Our results showed that the age-related changes in mRNA clustering differed by compartment (**Figure 5B**), implying that aging may drive compartment-specific functional modifications.

**Figure 5.**
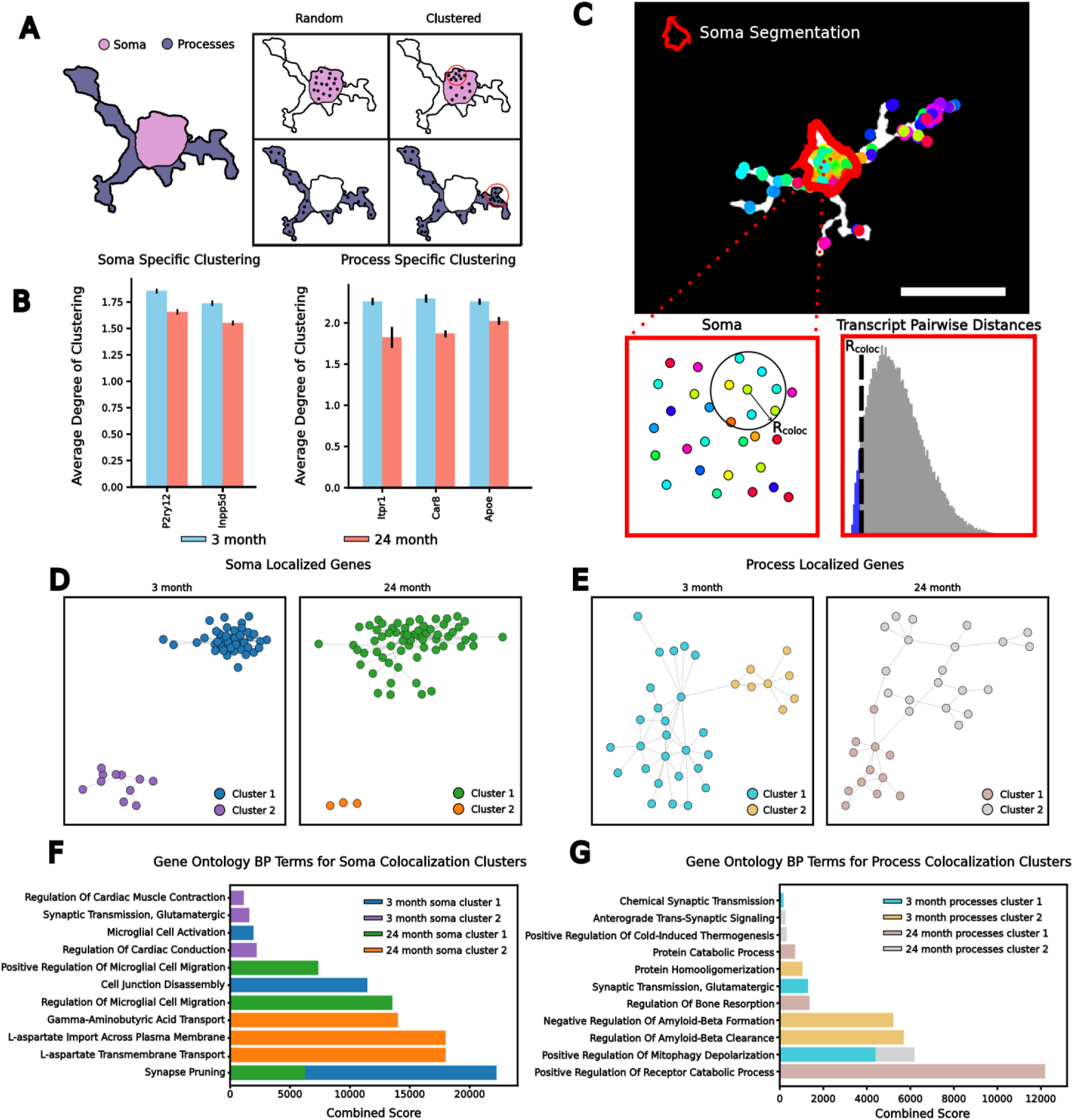
Compartment Specific Gene Colocalization Changes with Age. **A,** Schematic comparison between completely random spatial null distribution and spatially clustered sample for both the processes and the soma. **B,** Comparison of degree of clustering within each compartment with age, error bars show the standard error of the mean, and genes were only showed if deviation with age was statistically significant by the FDR adjusted p-values from the Wilcoxon-Rank Sum test. **C,** Schematic demonstrating identification of gene-gene pairs with a colocalization threshold R_coloc_ (**top**) and schematic for how th**e** R_coloc_ was set (**bottom**). Scale bar is 25 µm. **D**, Statistically significant soma-specific gene-gene localization networks for both young (left) and aged (right) microglia. The color in both graphs reflects gaussian mixture model clustering of the individual networks. **E**, Statistically significant process-specific gene-gene localization networks for both young (left) and aged (right) microglia. The networks are colored by gaussian mixture model clustering. **F**, Significant gene ontology biological processes across both young and old soma-specific gene localization networks. The colors reflect the spectral clustering colors defined in **D**. **G** Significant gene ontology biological processes across both young and old process-specific gene localization networks. The colors reflect the spectral clustering colors defined in **E.**

We hypothesized that such age-dependent shifts in mRNA clustering could affect function by altering networks of colocalized transcripts^64,65^. To test this, we first calculated pairwise distances between all transcripts localized to the soma and used the bottom fifth percentile of these distances as our colocalization threshold (**Figure 5C**). For each pair of genes, we quantified the number of corresponding transcripts that fell within this threshold. To evaluate whether the observed colocalization exceeded random expectations, we generated null distributions by randomizing RNA positions within the segmented soma and process compartments and recalculating colocalization across 1,000 randomized cells (**Methods**).

This approach revealed significant age-related reductions in mRNA colocalization. In the soma, 171 gene pairs showed significant colocalization in young microglia versus only 126 in aged microglia. Similarly, in the processes, 48 gene pairs were colocalized in young cells compared to just 37 in aged cells (**Figure 5D & 5E**; **Supplementary Sheet 2**). Clustering these gene networks (**Methods**; **Figure 5D, E**) identified functional gene sets that differed by both compartment and age (**Figure 5F, G**).

In the soma, younger mice (3-month-old) displayed differences in functional gene clusters than aged mice (24-month-old). These younger clusters were enriched for GO terms related to synapse pruning, cell junction disassembly, and an “activated” microglial state. In contrast, the aged soma network was associated with GO terms related to microglial migration (**Figure 5F**). Turning to the processes, we found that aging also reshaped the functional composition of colocalized gene networks. In young mice, process-enriched networks were associated with negative regulation of amyloid beta formation and synaptic transmission, whereas in aged mice, these networks were linked to mitochondrial depolarization and positive regulation of catabolism (**Figure 5G**). These findings align with previous studies demonstrating that age-related changes in mRNA colocalization can influence downstream brain functions^66^. Taken together, our results underscore how mRNA compartmentalization underlies age-associated shifts in microglial function, with distinct compartment-specific transcript interactions driving differential biological outcomes.

## Discussion

The spatial organization of transcripts within cells provides crucial insights into their functional states^67^, yet this relationship has remained largely unexplored in microglia. While characterization of microglial morphology through fluorescent imaging has long been the accepted standard for identifying “activated” cells in both subclinical and clinical models^68,69^, the connection between transcript localization and morphological states has been unclear. The paradigm that morphology is intertwined with biological function is fundamental to frameworks that infer the transcriptomic landscape of large tissue regions from histological images^70^. These inferences, although accurate on the macro scale, had not been investigated at the level of the individual cell or its subcellular compartments. In this study, we present an integrated framework using MERFISH paired with fluorescent immunohistochemistry to simultaneously analyze the morphological and transcriptomic landscapes of individual microglia, including the spatial distribution of transcripts within cellular compartments. This approach revealed heterogeneity in the morphologies of different transcriptomically defined microglia and linked these morphological changes to subcellular localization of specific gene subsets in the transcriptome. Furthermore, we revealed discrepancies between microglial morphological state and transcriptional state while uncovering age-related changes in mRNA compartmentalization. This allowed us to identify gene sets enriched within the microglial branches and use them to classify the ramification status of different cells. This study identified a transcriptomic signature that can be used to identify different morphological states underlying different transcriptomically defined microglial subtypes.

The long-standing doctrine has been that microglia “activation” is followed by a retraction of their branches. Although a consensus report in the field has begun to question this^44^, it remains the predominant view in microglial biology for many. By leveraging high-resolution imaging and transcriptomic profiling of individual microglia, we identified heterogeneity in microglial morphology across different transcriptomic subtypes. Notably, the glutamate transporter gene *Slc1a2*, whose transcripts predominantly localized to the branches of microglia, was upregulated in more ramified microglia compared to amoeboid cells. Interestingly, increased expression of GLT-1, the protein encoded by *Slc1a2*, has been linked to “activated” microglia^45^, directly challenging the perceived correlation between “activation” and amoeboid morphology. Moreover, nonfunctional variants of GLT-1 have been associated with the loss of microglial branching^71^, further validating our findings. Additionally, our data revealed that amoeboid microglia, compared to ramified microglia, do not express differentially regulated genes related to phagocytosis or synaptic clearance within individual transcriptomic classes. These findings reveal that transcript localization serves as a critical determinant of microglial function, independent of traditional morphological classifications, suggesting new therapeutic approaches for modulating microglial function in disease states.

Disruption of microglial morphology has not only been seen as an “activation marker but also a consequence of aging^25^. Aging induces both structural changes in the CNS and dysregulation in the functional compartmentalization of RNA^5^. In our study, we observed that young microglia had a greater number of process-localized genes, and different gene regulatory networks, compared to their aged counterparts. These changes in RNA compartmentalization may result from altered RNA trafficking mechanisms within the cell^72^, which could serve as a protective strategy against the age-related degradation of mRNA in the cell body^1,73^. However, our data revealed that the process-localized genes were effective at predicting morphology regardless of age. This finding suggests that while aging affects the breadth of transcript localization, core mechanisms governing cellular architecture remain preserved throughout the lifespan. Understanding these conserved pathways could provide crucial insights into maintaining cellular function despite age-related deterioration of RNA trafficking systems.

The rapid development of spatial transcriptomic technologies is pushing resolution limits below the single-cell level^74,75^, enabling unprecedented insights into the subcellular organization of mRNA across various tissues^76–78^. This advancing technology reveals how the precise positioning of transcripts within cellular compartments contributes to both morphological complexity and functional specialization. Pairing these technologies with advanced imaging modalities will be essential for understanding how subcellular transcript localization shapes cellular heterogeneity and drives functional complexity. By integrating full transcriptome measurements with imaging of morphologically complex phenomena, correlational maps, such as those generated in this study, can help uncover how disrupted transcript localization contributes to diseases that manifest at the structural level^79,80^. Our analysis methodology extends beyond traditional cellular categorization, enabling investigation of genetic correlations with any morphological or subcellular feature identifiable through fluorescent microscopy. This ability to map transcript distribution patterns while simultaneously capturing cellular architecture moves us closer to understanding how spatial organization of RNA orchestrates cellular function. As these technologies continue to evolve, the integration of subcellular transcript mapping with morphological analysis promises to revolutionize our approach to precision medicine, offering new therapeutic strategies based on both cellular structure and the spatial organization of gene expression.

## Methods

### MERFISH Analysis

#### MERFISH Panel Selection

Using a combination of single-nucleus RNA sequencing data with aging and neuroscience literature we were able to select genes for MERFISH. Our selection criteria for the single nucleus RNA sequencing data involved identifying cell-type-marker genes for each cell population using a one-vs-all approach. To do this we utilized an internal dataset with single nucleus sequencing of the cerebellum, and we utilized a previously published single nucleus dataset of the mouse hippocampus^50^. We then performed a Mann Whitney–Wilcoxon test for each gene between the cells within each cell population and all other cells not in that population and corrected the resulting P values for multiple hypothesis testing using the Benchamini-Hochberg correction to obtain false discovery rate-adjusted P values. As previously described^81^ a gene was considered a cell-type marker for a specific cell population if it: (1) was expressed in at least 30% of cells within a specific population; (2) the false discovery rate-adjusted P value < 0.001; (3) gene expression in the specified population was at least fourfold higher than the average expression in all cells not in that population; and (4) expressed in a fraction of cells within the specified population that was at least 2 times higher than any other population of cells. We then saved the top five marker genes for each cell type based on the effect size of the log fold change. In addition to these markers, known genes related to age^50^, microglia, astrocytes, oligodendrocytes, neuronal markers^81^, and genes related to subcellular structure from the literature were included which brought the panel to a total of 500 genes.

#### Tissue Processing for MERFISH

Brain samples for all mice were processed using a fixed-frozen protocol. After anaesthetization with 2.5% v/v Avertin, transcardial perfusion with 20 ml cold (4 °C) PBS was followed by perfusion with 30 ml of 4% paraformaldehyde, also maintained at 4 °C. The brains were immediately removed and submerged in 4% paramformaldehyde for an overnight incubation at 4 °C. Following incubation, the brains were then placed in 30% sucrose until sink. Samples were then hemisected and frozen in OCT using dry ice, after which they were stored at −80 °C until sectioning. Sectioning was performed along the sagital plane on a cryostat at −20 °C. Slices between 10 and 20 μm in thickness were captured onto Vizgen slides (Vizgen; REF 10500001) for MERFISH.

### Sample Preparation for MERFISH Imaging

Vizgen slides with single mouse brain sagittal sections were processed according to a fixed-frozen tissue sample preparation MERSCOPE protocol (Vizgen). The slides, upon removal from the cryostat, were washed three times with 1X PBS and placed in 70% ethanol at which point they were placed in a photobleacher (Vizgen) for 3 hours, following photobleaching the samples were stored at 4 °C overnight. Following incubation overnight, the samples were washed with 1X PBS and covered with Blocking Buffer C Premix (Vizgen; REF 20300100) and RNase inhibitor (NEB; REF M0314L) for one hour. The samples were then placed in a primary antibody staining solution consisting of Blocking Buffer C Premix, 10% RNase inhibitor, and IBA1 antibody (Abcam; REF ab178846) at a volume dilution of 1:1000 for ninety minutes. Primary antibody incubation was then followed by three washes with 1X PBS. Following washing the samples were then placed a secondary antibody staining solution consisting of Blocker Buffer C Premix, 10% RNase Inhibitor, and an Anti-Rabbit Aux 5 (Vizgen; REF 20300102) at a 1:100 dilution for one hour. The samples were then washed three times in 1X PBS followed by a 15-minute incubation with 4% paraformaldehyde and an additional two washes in 1X PBS. Following the staining protocol the samples were then washed one time with Sample Prep Wash Buffer (Vizgen; REF 20300001), then washed with Formamide Wash Buffer (Vizgen; REF 20300002) for 30 minutes at 37 °C. A 500 gene panel mix (Vizgen; REF 10400003) was then incubated on the tissue samples for anywhere between 36 and 48 hours at 37 °C. After hybridization, the samples were washed with Formamide Wash Buffer 2 times at 47 °C for 30 minutes each. The tissue samples were then embedded in a 4% polacrylamide gel and were treated using the Clearing Premix (Vizgen; REF 20300003) and Proteinase K (NEB; REF NC0547027) overnight at 37 °C to digest proteins and lipids in the samples. Following the digestion the coverslips were washed two times with Sample Prep Wash Buffer, stained with a DAPI/PolyT mix for 15 minutes, and washed with Formamide Wash Buffer followed by Sample Prep Wash Buffer one final time each before imaging. Finally, the slides were loaded into the MERSCOPE Flow Chamber and imaged at 20x magnification to identify regions of interest and 63x magnification to image the probes.

### MERFISH Data Processing and Quality Control

MERFISH imaging data was processed with the MERlin^82^ pipeline to decode the spatial location of the mRNA probes. The cells were segmented using Baysor 0.6.2^34^ with the scale parameter set to 6.5 microns and the minimum molecules per cell parameter set to 50. The decoded molecules were then assigned to the cell boundaries from Baysor to produce a cell-by-gene matrix that lists the number of each molecules decoded for every gene within each cell. Each cell in the matrices was then filtered based on quality control cutoffs of minimum twenty transcripts per cell and minimum 5 unique genes per cell. The expression matrix for each sample was then concatenated and normalized and log transformed using Scanpy^83^ prior to integration with Harmony^84^. The samples were then subject to label transfer from the Allen Brain Cell Atlas^85^, following the Seurat Integration and label transfer pipeline^28^, and from an atlas of the adolescent mouse brain (http://mousebrain.org/adolescent/downloads.html) with the scVI and scANVI protocol^86^.

### IBA1 Initial Segmentations

The microglial IBA1 stain was imaged across 6-7 imaging planes, spaced by 1.5 microns each, using an anti-rabbit oligonucleotide conjugated antibody. Python’s scikit-image library was used to create a max projection image for each imaging stack. Following max projection, the images were preprocessed for segmentation following histogram equalization by being converted down to 8-bit images. Then, using OpenCV a background subtraction and edge detection was applied to the 8-bit images, to identify the borders of the microglia and remove any imaging noise caused by tissue autofluorescence. Finally, a Gaussian blur and an Otsu segmentation was applied to the images to generate coarse-outlined microglia. This binary image was then converted into a label image such that every segmented IBA1 stain was assigned an identity for downstream processing.

### Matching Transcriptomically-Derived Boundaries to Cell Masks

Each brain has a Baysor-derived cell segmentation associated with it, using these geometries we then align our labeled IBA1 image to cell segmentations. This allowed us to generate a list of cell boundaries that were derived transcriptomically and overlapped with the image analysis-derived cell boundaries. Then utilizing cell annotations which were generated from the Seurat based label transfer with the Allen Brain Cell Atlas, and cross-referenced against cells labeled using scVI and scANVI, we identified microglial labeled cell boundaries and binary image labels which were uniquely mapped to one another. The coordinates of these uniquely mapped microglia were then saved for further analysis.

### High-Dimensional Embedding of Microglial Morphology

Utilizing the microglial coordinates and coarse-grained morphology masks for each cell a bounding box was generated around every microglia in their raw max-projected image. Then following previously established methodologies for high-dimensional image embeddings^37^ we resized our microglia bounding boxes to 112x112 pixels and applied padding with a middle gray color to achieve a 224x224 image for each cell. These resized images were then passed through the VGG-19 neural network with pre-trained weights from the ImageNet dataset, as accessed through the pytorch Keras library. After passing the resized images through the neural network we generated a max-pooled vector of the fifth activation layer, to create a 512-dimension vector to represent the morphological space of the microglia.

The UMAP visualization and clustering, which were used to represent the morphology space was generated by performing principal component analysis on the 512-dimension vector. To generate the UMAP a nearest neighbor mapping of the top 10 principal components, capturing approximately 75% of the cumulative explained variance, was used for its calculation. The clustering which was used to represent the morphology space was calculated using the python kneed library and kmeans clustering. The kneed library’s KneeLocator function was used to identify the elbow of the kmeans cluster to inertia plot. Setting the number of clusters to the identified elbow we generated a clustering that captured the minimum amount of variance necessary to explain the data.

### Remapping Transcriptomic Data to Newly Segmented Microglia

Wanting to create a more fine-grained representation of the microglial transcriptome than what the Baysor segmentations offered we utilized the microglial coordinates to remap transcripts that were directly correlated with the microglial processes. To perform this analysis, we identified the IBA1 stain which was at the center of the saved microglial coordinates and performed an additional round of segmentation on the raw max projected microglia image and DAPI image. Utilizing OpenCV frameworks for adaptive thresholding, with a ∼200-pixel window size, we generated separate single-cell masks for the DAPI and the IBA1 channel aligned to each microglial coordinate. We then utilized the differences between the DAPI and IBA1 channel to generate a ‘soma’ image mask and a ‘processes’ image mask, with the unity between the two channels being our total cell mask. Applying the decoded transcript location outputs from MERFISH imaging runs we were able to correlate our soma, process, and total imaging masks to transcripts across all z-planes. These separate correlations were saved for each cell, rendering a soma, non-soma, and total count vector for each.

### Annotation of Transcriptomically Defined Microglial Clusters

The total count for each microglia was normalized by dividing each gene’s total transcript count by a cell’s total amount of transcripts and multiplying by 10,000, then ultimately log transforming the data. The transcriptomic UMAP visualization was created by scaling the log-transformed counts and calculating principal components on all the genes in the dataset. The top 40 principal components were utilized to generate the nearest neighbor graph for UMAP calculation, and leiden clustering of the principal components was used to sub-cluster the microglia. Individual subclusters were tested for marker gene expression based on the Wilcoxon-Rank Sum test, to aid in the identification of sub-cluster identity. Sub-clustering identification was ultimately concluded based on manual annotation from the microglial subtype marker gene expression, normalized between the different leiden clusters.

### Microglial Morphological Measurements

Utilizing the same re-segmentation approach applied earlier we wanted to generate detailed morphological measurements of the microglial ramifications and features that have been described previously^40–43^. The individual characteristics were calculated by generating a region property for each microglial segmentation and utilizing native skcikit-image libraries to calculate the features. In total, pixelwise morphological measurements were calculated for cell area and perimeter, the convex hull area and perimeter, solidity, convexity, roughness, circularity, convex hull’s span ratio, convex hull circularity, eccentricity, the Euler number, the extent, soma area, soma perimeter, soma circularity, soma to cell size ratio, and the cell’s ferret diameter. The next set of measurements were a direct result of skeletonization of the microglial segmentation in which we analyzed the total skeleton length in pixels, the mean length of each branch, the total number of branches, the total number of branching points, and the number of terminal points, we then calculated Euclidean and path distance of the skeleton branches. We then assessed sholl parameters by calculating the number of intersections between the skeleton branches with circles of varying radii centered around the central node. The sholl parameters calculated were the ramification index, which is a ratio between radius of the circle with the greatest number of intersections and the radius of the first non-zero circle, the radius with the greatest number of intersections, and the largest radius which had an intersection with the skeleton. Fractal dimension^41^, lacunarity^41^, and tortuosity^87^ were also calculated for the individual microglia. Finally, we assessed the normalized intensity of the IBA1 stain for each of the region properties assessed.

### Comparison Between Differentially Expressed Genes from Transcriptomically and Morphologically Derived Microglia

To compare microglial labels which were derived transcriptomically against those that were derived morphologically we performed differential expression analysis which was estimated using the Mann–Whitney–Wilcoxon test. We compared the DAM microglia to the Homeostatic microglia, and then we compared the C1 morphology cluster to the C5 morphology cluster. We categorized a differentially expressed gene as one that displayed a log2 fold change greater than 0.5 and an unadjusted p-value of less than 0.05. Following the identification of differentially expressed genes we identified enriched Gene Ontology Biological Process terms for each set of genes, morphology or transcriptomic derived clusters, using the Python gseapy package. To compare the glutamate transporter score between C1 and C5 microglia in the homeostatic cells we scored C1 and C5 cells for log-normalized expression of *Slc1a2*, *Gria2*, *Dlgap2*, *Dlg2*, and *Dagla* ^45,88–92^ using scanpy’s tl.score_genes functionality and performed a Mann–Whitney– Wilcoxon test between the two distributions to establish statistical significance.

### Correlation Between Morphological Measurements and Transcript Measurements

We utilized the MERFISH and morphological data for each individual microglia in this analysis. Following log normalization of the total transcript counts for each cell, we then performed a training-test split with 75% of the cells in our object being used for training and 25% of the cells being used for following clustering analysis. Using the cells from our training set, we calculated the spearman correlation coefficient for each morphology-gene pair. Generating a list of the genes which were amongst the top 5 correlating for each individual morphological feature, we then clustered the microglial morphological clusters in our test set using the top 5 correlating genes. This analysis was then repeated on each individual brain region across the mice. Wanting to better understand the correlation between the genes and the features that correlate with higher degrees of ramification we selected the genes with the highest degree of correlation to the following features: ‘Cell Area’, ‘Convex Hull Perimeter’, ‘Convex Hull Area’, ‘Cell Perimeter’, ‘Number of Branches’, ‘Skeleton Length’, ‘Cell Roughness’, ‘Number of Terminal Points’, ‘Number of Branching Points’, ‘Soma Ratio’, ‘Path Distance’, ‘Fractal Dimension’, ‘Euclidean Distance’, ‘Tortuosity’, ‘Branching Index’, ‘Dendritic Maximum’. The genes selected were generated from observation of the cluster map created from displaying correlations between the top 5 correlating genes for each morphology feature, and these genes included: *Slc1a2, Gria2, Pink1, Ank2, Gria1, Slc6a11, Aldoc, Gnas, Mfn2, Atp2a2, Mobp, Pcdh9, Gfap, Aqp4, Sdc3, Cisd1, Sst, Spp1, Apoe, Axl, Pcdhb8, Cxcl3, Rbm3, Cemip2, Cirbp*, and *P2ry12*. These gene and morphology sets were then calculated for average spearman correlation in each of the individual regions, where the average was for each gene across correlations for all morphologies. This was used to generate a count for how many regions each gene was a top 5 correlating gene for.

### Intracellular Clustering of Transcripts

We first calculated a normalized distance for every transcript from the center of the soma in our C4 and C5 microglia. To do so, for every C4 and C5 microglia in the dataset we visualized the soma, using a DAPI stain overlayed with the microglia stain, and used the regionprops.centroid function in scikit-image to establish a soma center. We then calculated Euclidean distance from the outermost edge of the microglia and every transcript to the centroid. We then normalized the transcript-centroid distances by the outermost edge-centroid distance for each microglia. These results were then visualized as a cumulative density function for each transcript species.

The degree of clustering was calculated from an adaptation of the DypFISH Ripley-K generalization^63^. In brief we estimated a normalized Ripley K function for each transcript under a homogenous Poisson process to create a Ripley H function for each cellular compartment. We then compared the observed Ripley H function to a compartment-specific Ripley H function derived from a random distribution of transcripts across the whole cell. Spatial clustering was considered significant at a radius r if the computed Ripley’s K is above the 95% or below the 5% confidence interval for the random distribution. We performed this analysis in each of the two compartments individually, the soma clustering was performed only on the transcripts which overlapped with the underlying DAPI stain. The processes degree of clustering was approximated by calculating the degree of clustering for a gene within each of the individual processes and averaging their values together. We then compared the significantly clustered gene’s degree of clustering values at each aging time point using the Wilcoxon Rank Sum test and only visualized genes with an adjusted p-value of less than 0.05.

### Comparison of Compartment Specific Gene Colocalization

For each C4 and C5 microglia in the dataset we calculated the pairwise distance between every transcript within the soma. Then by using the numpy.percentile function across all of the calculated distances we found the fifth percentile of one micron to be the colocalization radius. We used the subcellular location of each transcript to identify gene-gene pairs that were within one micron of each other. We also generated a null distribution of transcript colocalization by randomizing the position of the transcripts within each cell and counting the gene-gene pairs separated by less than one micron. Similar to previous methods utilized for cell-cell interaction^28^ we randomized and repeated this process 1000 times to generate a null distribution for each cell in the dataset. We then compared the observed number of gene-gene contacts to the null distribution and performed a Z-test with Benjamini-Hochberg correction. We calculate the threshold for significant colocalized counts by calculating the 97^th^ percentile of non-zero gene-gene contacts across our average null distributions for each cell. From this analysis we considered a gene-gene pair to be colocalized if it had 5 or greater observations, an adjusted p-value of less than 0.05, and formed a cluster of greater than 2 genes.

Following establishment of gene-gene pairs we subset our dataset down to just look at the C4 and C5 microglia. We generated networks of genes using python’s networkx library, with the edges being weighted by the z-score between the observed and the null distribution. We then apply scikit-learn’s GaussianMixture, with manually determined cluster numbers for each connected component to separate gene interaction networks into functional clusters. The genes comprising these clusters were then subjected to gene ontology analysis for biological processes.

### Comparison of Soma Enriched vs Process Enriched Gene Expression and Correlation

To compare the enrichment of certain genes in the different intracellular compartments we performed differential expression with the Mann–Whitney–Wilcoxon test. This differential expression was performed on only the most ramified microglial set, those microglia mapping to ‘C4’ and ‘C5’ morphologies so that we maximized the process area in comparison to the soma. Then independently taking the soma and non-soma counts and log-normalizing the two sets as described previously for the same set of cell we performed the Mann–Whitney–Wilcoxon test between them. We described process-enriched genes as those which held a log2 fold change of one with an adjusted p-value of less than 0.05 when compared to the soma-enriched genes, and the same criteria was applied to the soma-enriched genes. This analysis was repeated for the whole data set, regardless of age, and for the dataset split by age.

Following the selection of process- and soma-enriched genes for each age in the data set we took the genes which were the intersection of the old and young process and soma-enriched genes. This yielded one set of process-enriched genes and one set of soma-enriched genes. We then calculated the spearman correlation coefficient for each of the individual genes were every morphological characteristic regardless of age. We then averaged coefficients for each morphology characteristic for each set of enriched genes. This analysis was then repeated for the branch-enriched genes at each age.

### Random Forest Classifier Creation and Performance Analysis

To assess the predictive capacity of the branch-enriched genes as a whole and at each age we wanted to establish a classifier for the different morphology classes. To build this classifier, we selected the universal branch-enriched genes as our features. Then we categorized morphologies C1 and C2 together as amoeboid and we categorized C4 and C5 together as ramified and generated a model to predict if the cells were amoeboid or ramified in general and for each age.

To train the random forest models, utilizing scikit-learn’s train_test_split function we generated and 80/20 training test split for the microglia both as a whole and at each age. Then for training we utilized scikit-learn’s GridSearchCV to tune the hyperparameters of the random forest classifiers to achieve greatest accuracy for each. Assessing the performance of these models we utilized the AUROC, calculated through scikit-learn’s roc_auc_score. Utilizing the mean decrease in impurity calculation for each of the features which were used to train the classifier we assign feature importances to each gene as a whole and for the individual ages.

## Supporting information

Supplemental Sheet 1

Supplemental Sheet 2

## Data Availability

Decoded MERFISH data and Baysor processed cell boundaries collected during this work are available on figshare (https://figshare.com/articles/dataset/Aging_MERFISH_Brains/27919227). The images used to decode cell position and visualize microglia are available upon reasonable request.

## Code Availability

All computational analyses were performed using Python 3.9 and we performed cell segmentation using Baysor 0.6.2. All code for the work and the manuscript are available on GitHub at https://github.com/doughenze/Microglial_Morphology. The environments used to run the analysis are stored as singularity containers on Zenodo at: https://zenodo.org/records/14611122.

## Acknowledgments

The authors thank all members of the Quake Lab for valuable feedback and discussions throughout the study. The authors would also like to thank the members of the Wyss-Coray Lab and the Chan Zuckerberg Biohub for their support throughout the study. The authors also acknowledge all of the animals sacrificed for this study.

## Author’s Contribution

D.E.H and S.R.Q conceptualized and designed experiments. D.E.H and A.P.T collected data and conducted experiments. D.E.H analyzed the data. D.E.H and S.R.Q discussed and interpreted the data. D.E.H designed the figures. D.E.H and S.R.Q wrote the manuscript. D.E.H, A.P.T, T.W.C, and S.R.Q supervised, directed, and managed the study. All authors discussed the results and commented on the manuscript.

## Competing Interest

The authors declare no competing interests.

## Extended Data Figures

**Extended Data Figure 1:**
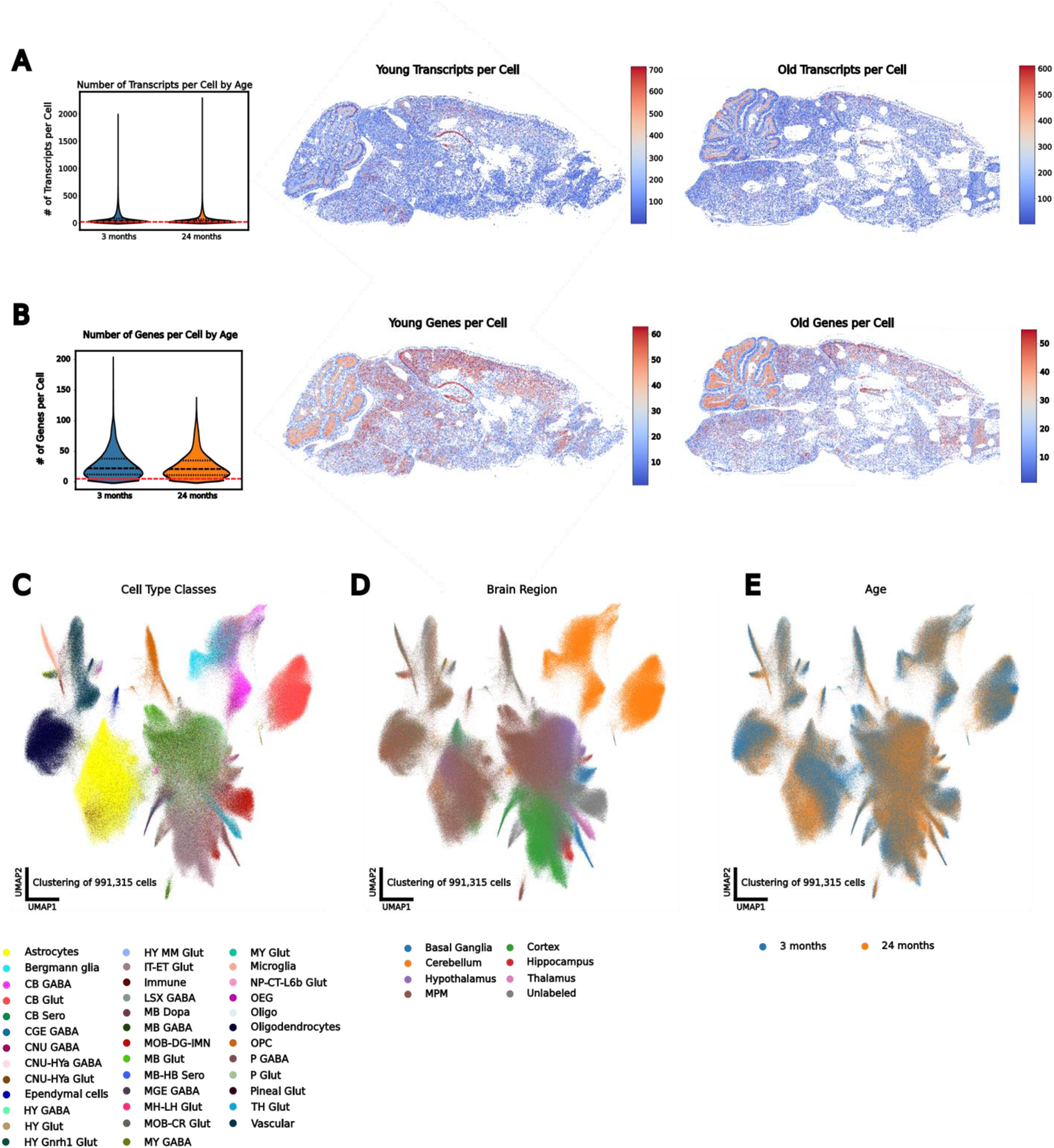
Spatial Transcriptomics Resolves Cell Types Across the Aging Mouse Brain. **A,** Number of cells passing minimum number of transcripts per cell threshold (red line) (left) with a representative spatial heatmap in a young (middle) and old (right) brain. **B,** Number of cells passing minimum number of genes threshold (red line) (left) with a representative spatial heatmap in a young (middle) and old (right) brain. **C,** UMAP representation of the spatial transcriptome of all cells passing QC, colored by major cell class. **D,** UMAP representation of all cells passing QC, colored by brain region. **E,** UMAP representation of all cells passing QC, colored by age.

**Extended Data Figure 2:**
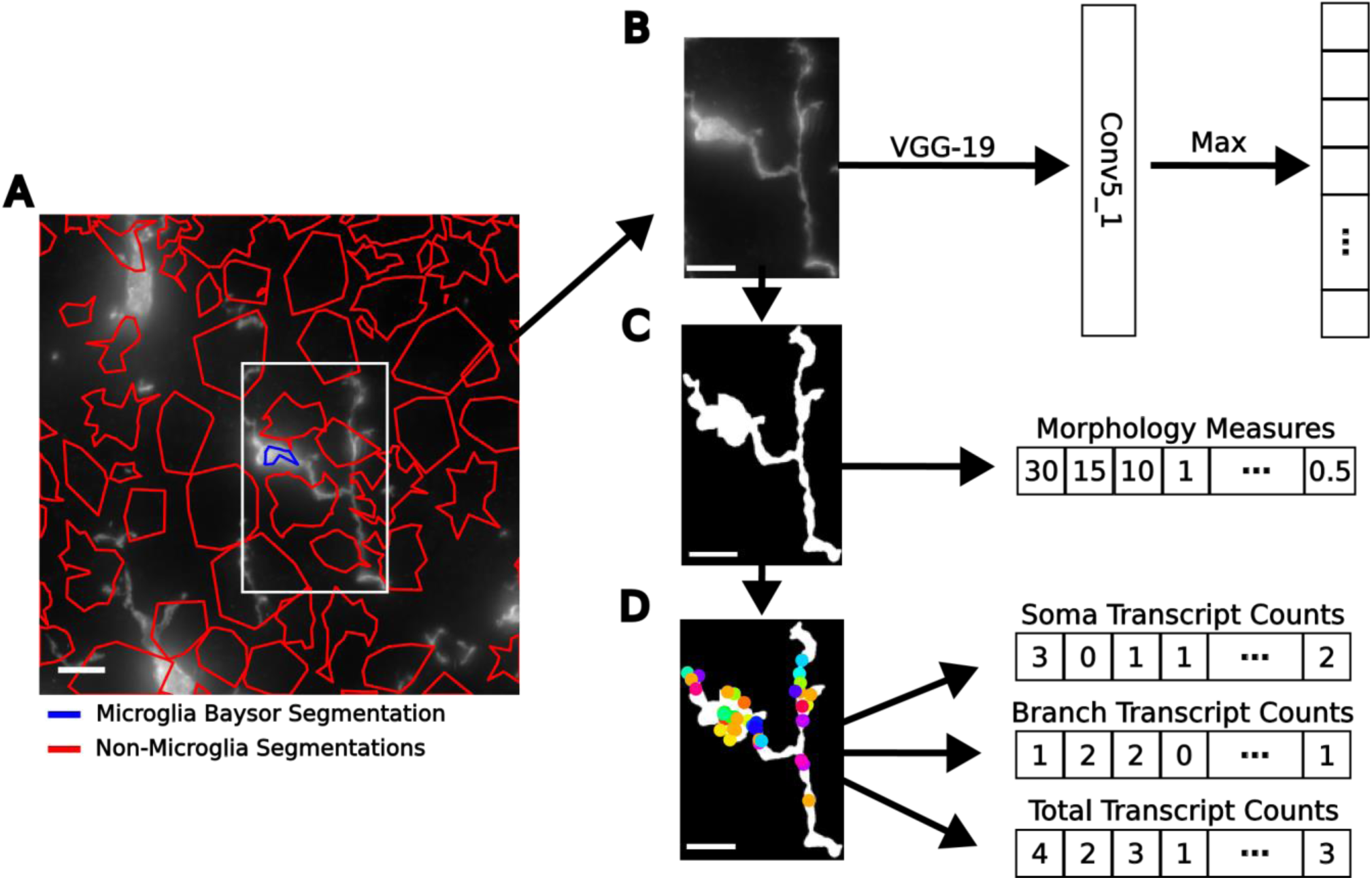
Schematic of Analysis Pipeline. **A,** Matched Baysor cell segmentations with microglia-stained image, colored by selected (blue) and non-selected (red) cells. **B,** Morphological embedding strategy for raw microglial single cell image. **C,** Representative microglial cell segmentation and schematic representation of manual analysis yielding a morphology vector. **D,** Decoded transcripts overlayed onto microglia morphological rendering, and schematic representation of three separate counts tables generated for each cell. Scale bars are 10 µm.

**Extended Data Figure 3:**
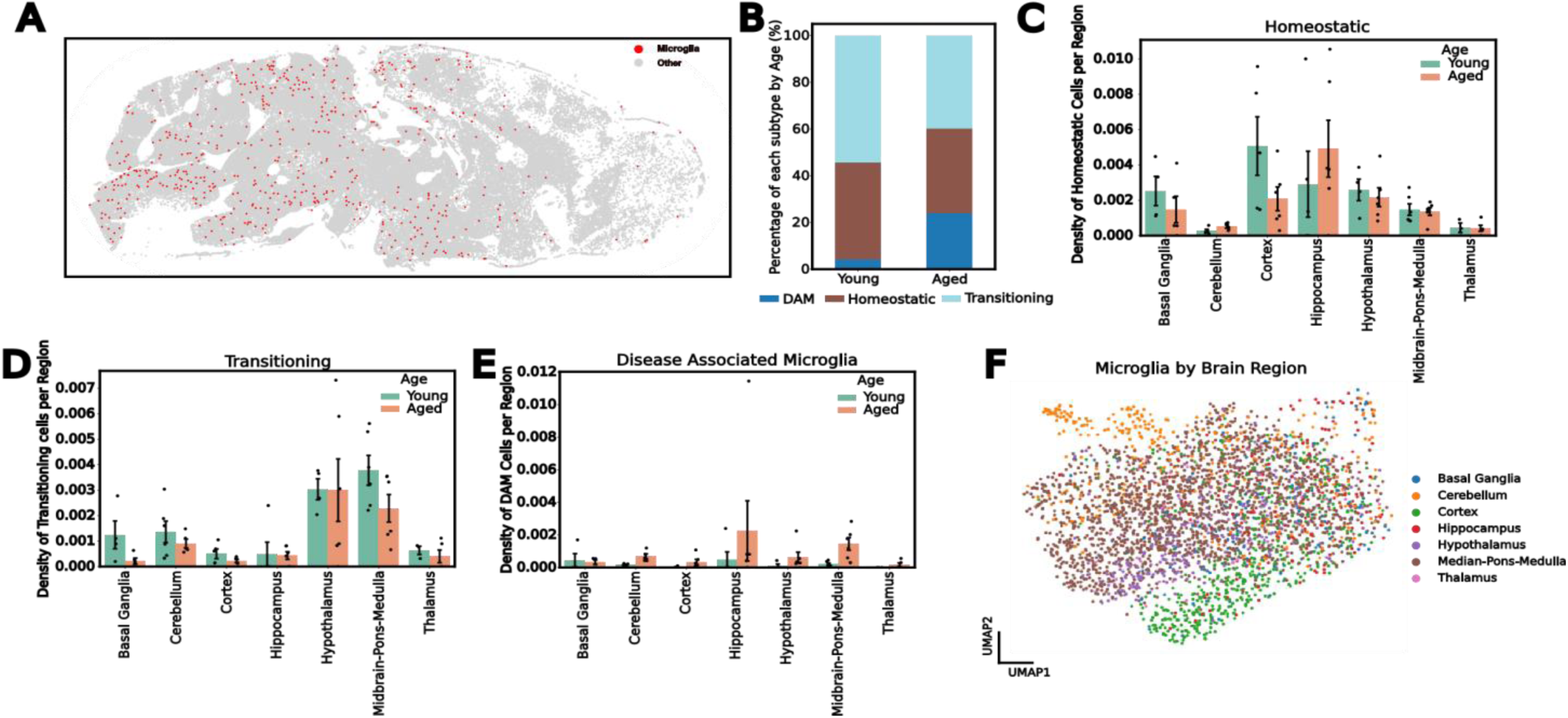
Spatiotemporal Patterning of Transcriptomically Defined Microglia. **A,** Representation of microglia distribution in the sagittal section of a brain. **B,** Frequency of microglial transcriptomic clusters in each age group. **C-E,** Quantification of different microglial transcriptomic clusters distribution across the brain with age, including the Transitioning (**C**), the Homeostatic (**D**), and the DAM (**E**) defined microglia. **F,** Transcriptomic UMAP plots colored by brain region.

**Extended Data Figure 4:**
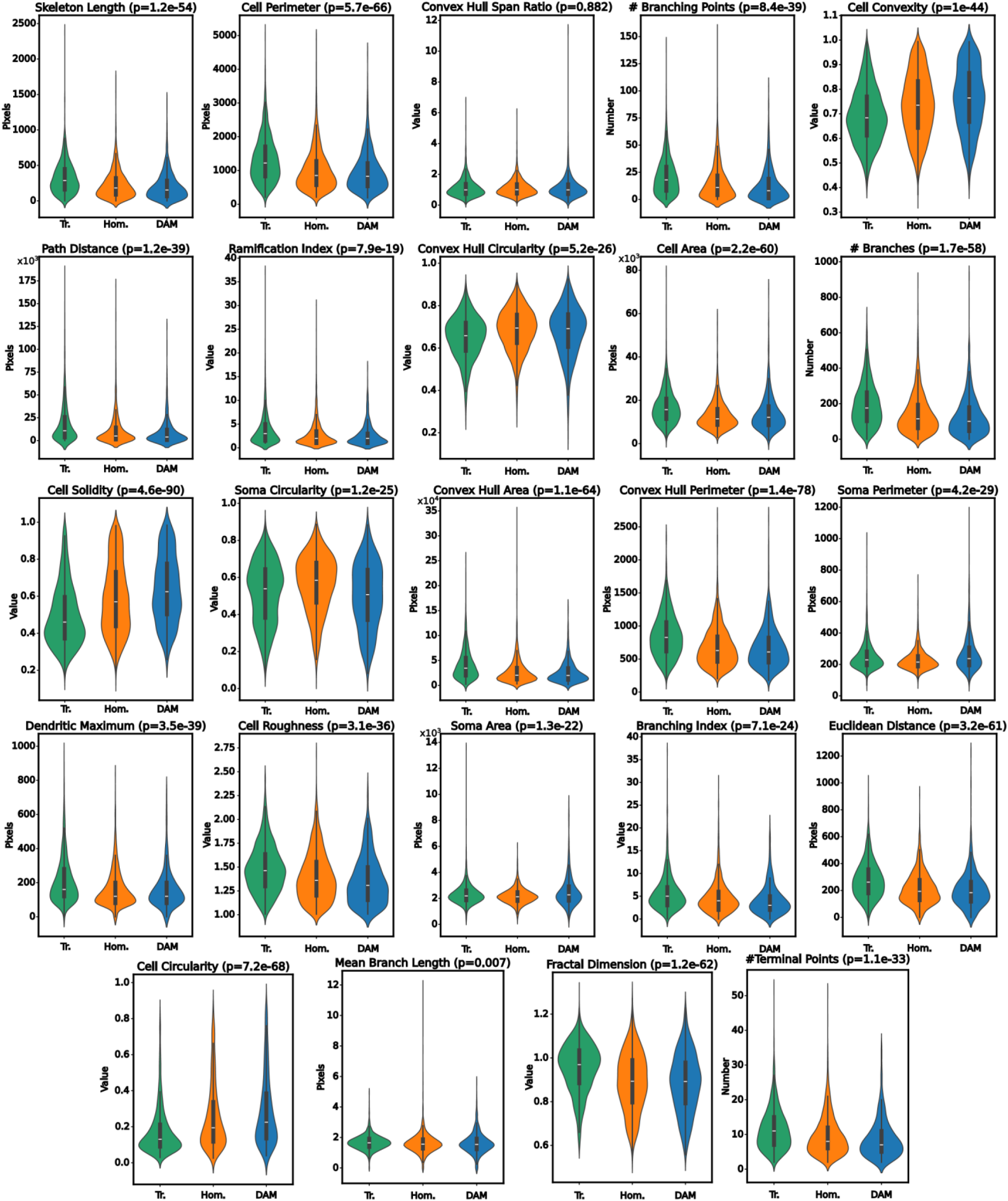
Morphological Characteristics of Transcriptomically Defined Microglia. Selection of quantifications of microglial morphological features, split by transcriptomic cluster. All measurements are either dimensionless, counting the occurrence of a phenomenon, or are pixel-wise quantifications. Each p-value was generated from an adjusted ANOVA.

**Extended Data Figure 5:**
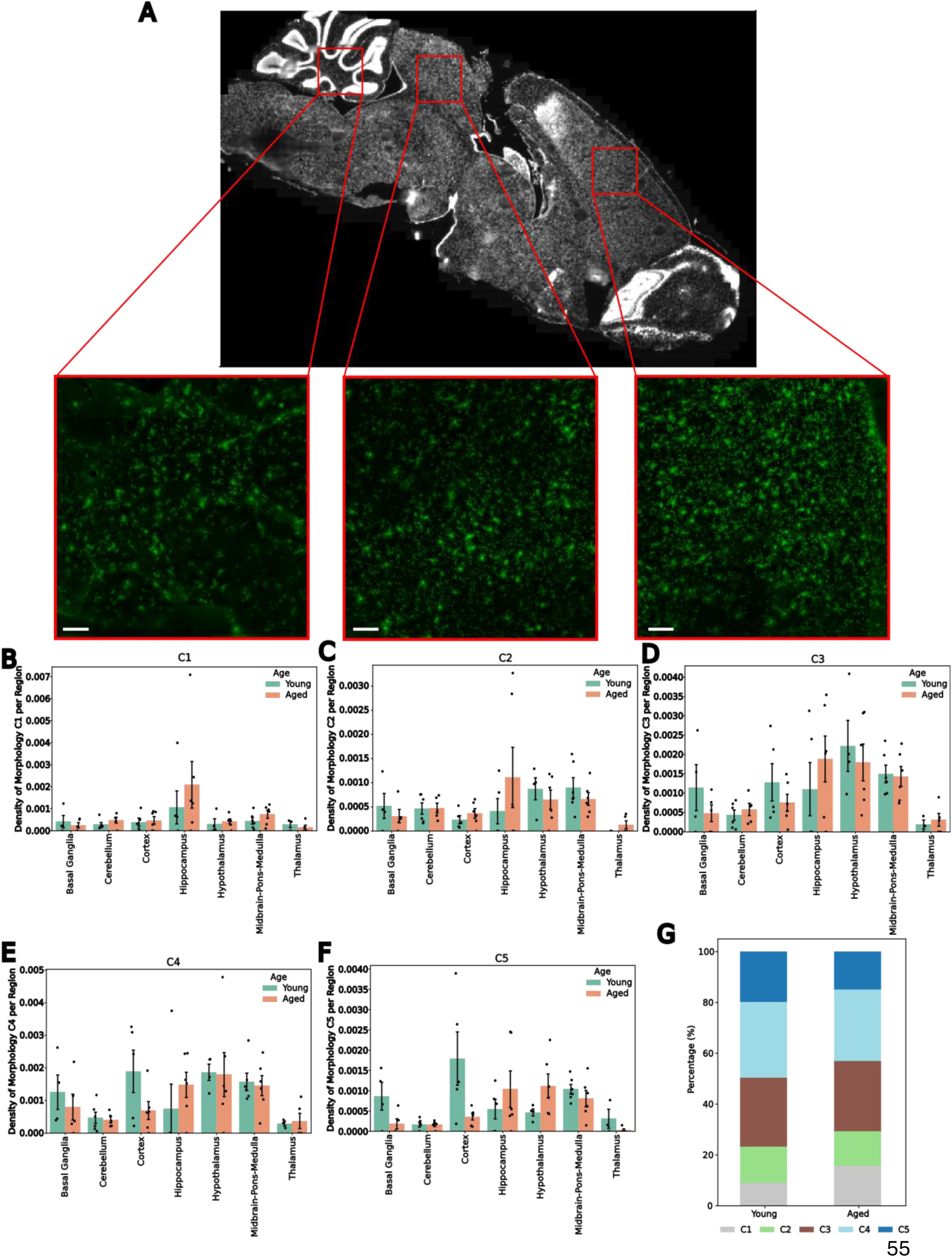
Spatiotemporal Patterning of Morphologically Defined Microglia. **A,** Schematic representation depicting IBA1 stained images as distributed in different regions of the brain. Scale bars are 100 µm. **B-F,** Quantification of different microglial morphological clusters’ distribution across the brain with age, including the amoeboid clusters C1 (**D**) and C2 (**B**) and the ramified clusters C3 (**C**), C4 (**E**), and C5 (**F**). **G,** Frequency of microglial morphological clusters in each age group. The error bars are standard error of the mean.

**Extended Data Figure 6:**
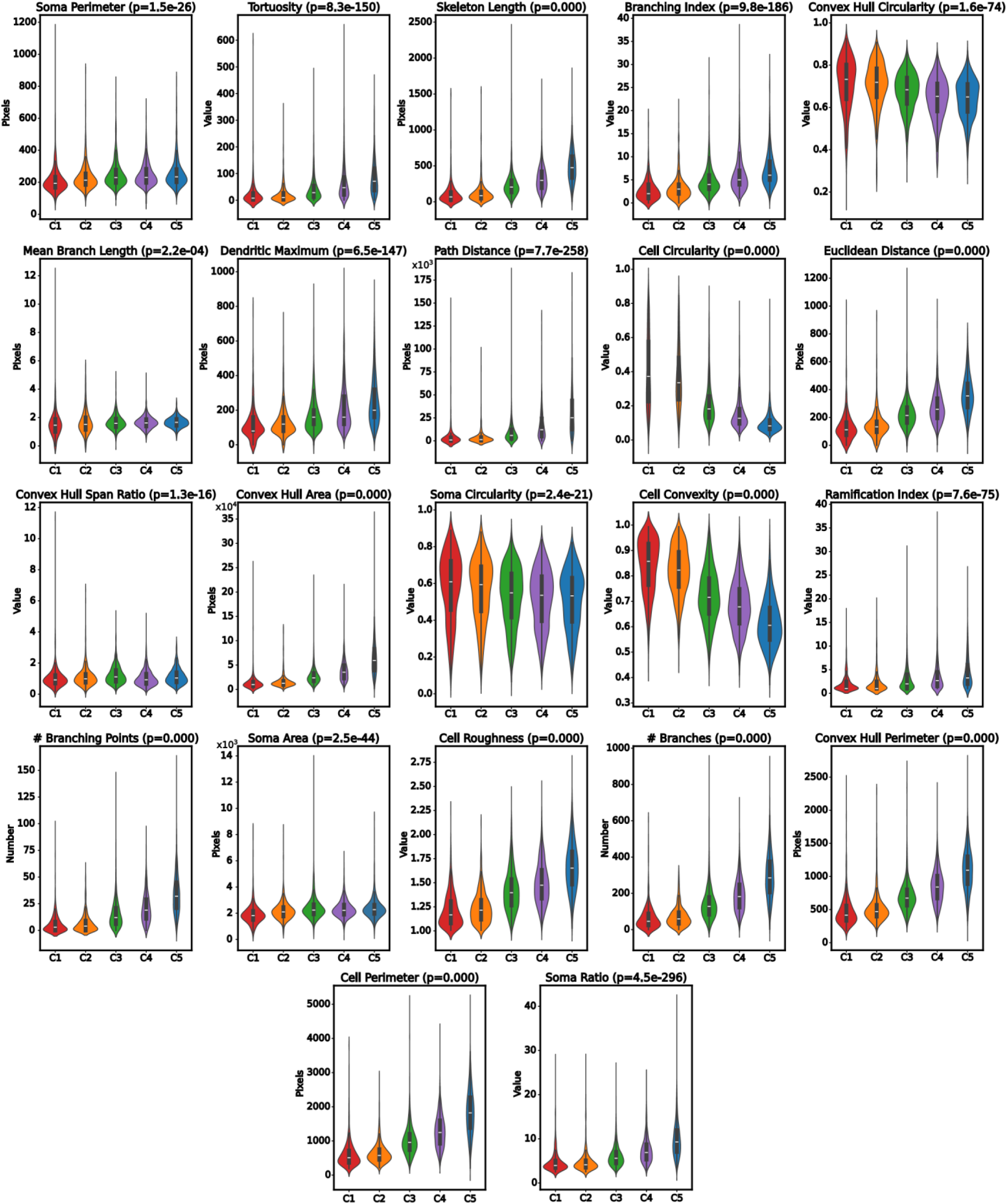
Morphological Features of Morphologically Defined Microglia. Selection of quantifications of microglial morphological features that did not appear in the main text, split by morphological cluster. All measurements are either dimensionless, counting the occurence of a phenomenon, or are pixel-wise quantifications. The p-values were generated from an adjusted ANOVA.

**Extended Data Figure 7:**
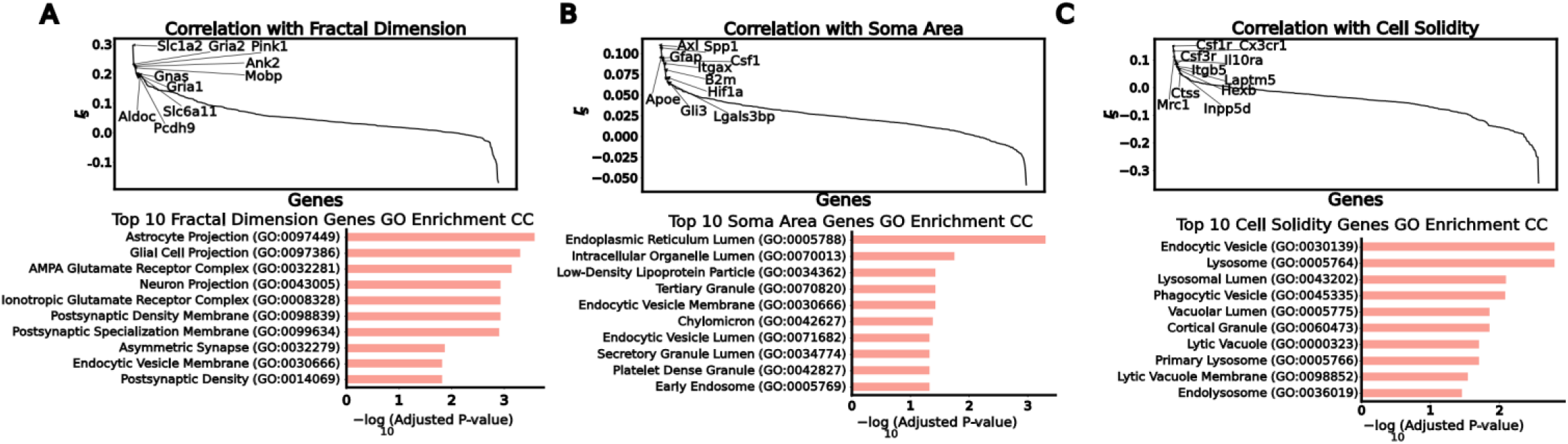
Single Morphological Characteristics Correlate to Compartment Relevant Genes A-C, Rank plots of Spearman correlation between selected morphological features and genes, top 10 genes for each feature are listed (top) and gene ontology cell compartment enrichment for the top 10 correlating genes for each morphological feature. The features selected include the Fractal Dimension **(A)** the Soma Area **(B)** and the Cell Solidity **(C).**

**Extended Data Figure 8:**
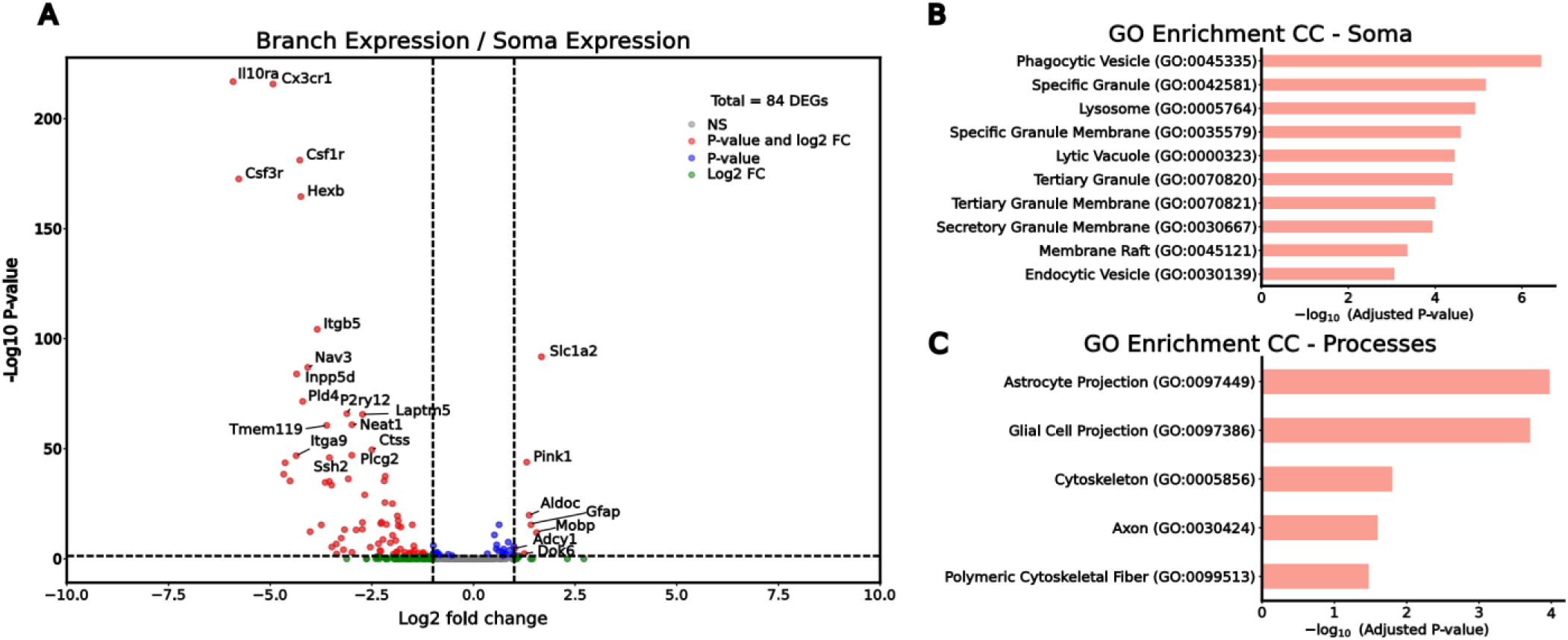
Soma and Branch Specific Transcripts are Enriched for Compartment Relevant Genes. **A,** Quantification of compartment enriched for each mRNA species in the panel. P < 0.05, adjusted P value by Mann–Whitney–Wilcoxon test. FC, fold change; NS, not significant. **B,** Gene ontology terms for cell compartment, calculated from the genes which were enriched in the soma. **C,** The same plot as in B, but the gene ontology terms were calculated from the branch enriched genes.

**Extended Data Figure 9:**
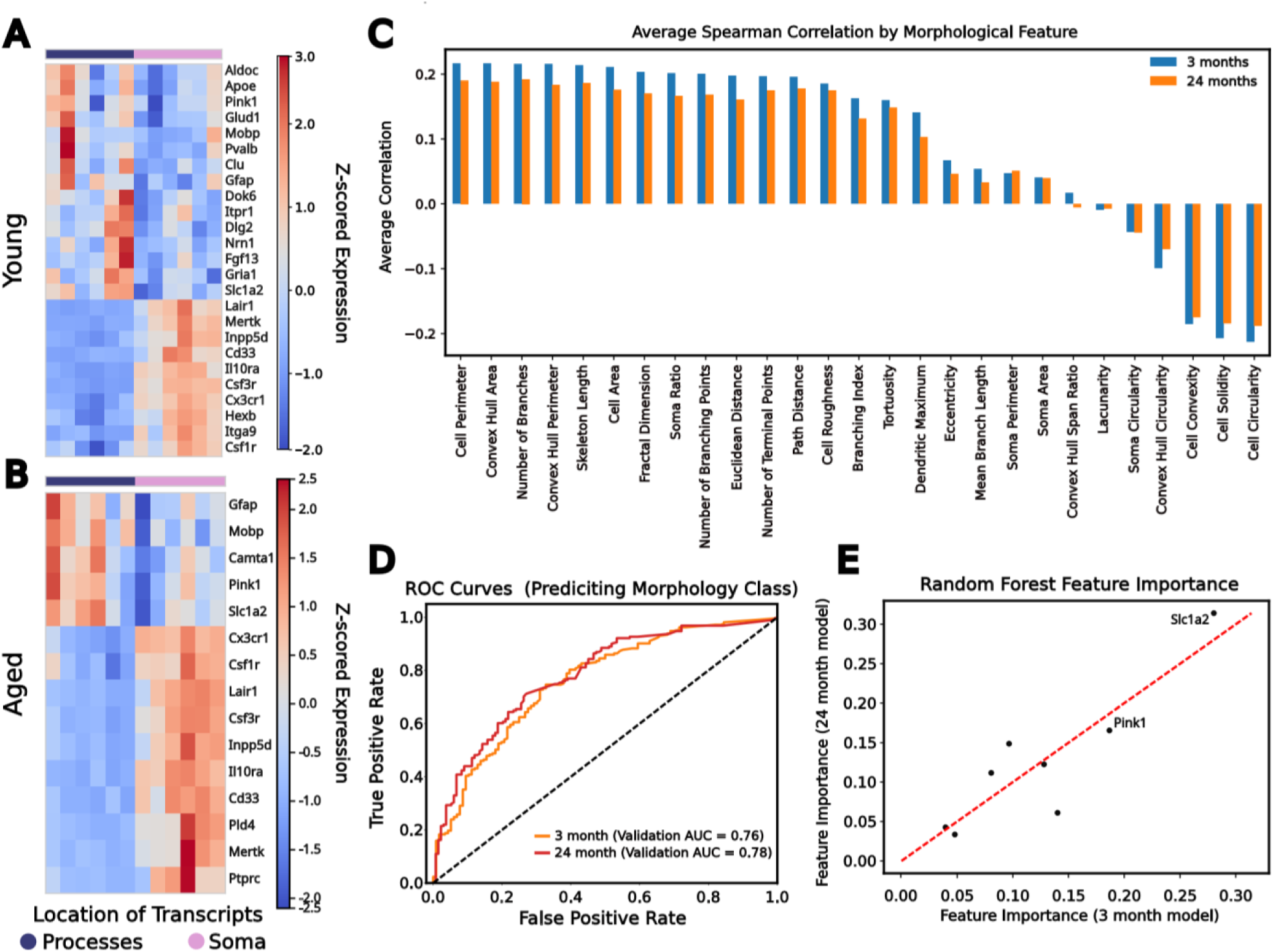
Compartment Specific Genes Are Affected by Age. **A,** Heatmap showing genes cluster by location of transcripts in young mice. **B,** Heatmap with aged specific transcripts clustering old mice by expression. **C,** Average correlation between process enriched genes with different morphological characteristics at each age, colored by age of mouse. **D,** Classifier performance as quantified by receiver operator characteristic curve (ROC) for samples from young or old mice. **E,** Importance of individual genes in each age stratified classifier as calculated through mean decrease in impurity.

## Notes

https://figshare.com/articles/dataset/Aging_MERFISH_Brains/27919227

### Competing Interest Statement

The authors have declared no competing interest.

